# Targeting the SUMO pathway primes all-*trans*-retinoic acid-induced differentiation of non promyelocytic Acute Myeloid Leukemias

**DOI:** 10.1101/254946

**Authors:** Hayeon Baik, Mathias Boulanger, Mohsen Hosseini, Julie Kowalczyk, Sonia Zaghdoudi, Tamara Salem, Jean-Emmanuel Sarry, Yosr Hicheri, Guillaume Cartron, Marc Piechaczyk, Guillaume Bossis

## Abstract

Differentiation therapies using All-*trans*-retinoic acid (ATRA) are highly efficient at treating Acute Promyelocytic Leukemia (APL), a minor subtype of Acute Myeloid Leukemias (AML). However, their efficacy, if any, is very limited in the case of non-APL AMLs. We report here that the inhibition of SUMOylation, a post-translational modification related to ubiquitinylation, restores the pro-differentiation and anti-proliferative activities of retinoids in non-APL AMLs. Controlled inhibition of SUMOylation with pharmacological inhibitors (2-D08 or anacardic acid), or *via* overexpression of SENP desumoylases, strongly enhances the ATRA-induced expression of key genes involved in differentiation, proliferation and apoptosis in non-APL AML cells. This activates ATRA-induced terminal myeloid differentiation and reduces cell proliferation and viability, including in AML cells resistant to chemotherapeutic drugs. Conversely, enhancement of SUMOylation by overexpressing the SUMO-conjugating enzyme Ubc9 dampens the expression of ATRA-responsive genes and prevents differentiation. Thus, inhibition of the SUMO pathway is a promising strategy to sensitize non-APL AML patients to retinoids and improve the treatment of this poor prognosis cancer, which has not significantly changed over the past 40 years.

## Introduction

Acute Myeloid Leukemias (AML) are a heterogeneous group of severe hematological malignancies. They arise through the acquisition of oncogenic mutations by hematopoietic stem- or progenitor cells. Instead of differentiating into normal blood constituents, leukemic cells are blocked at intermediate differentiation stages, proliferate and infiltrate the bone marrow, leading to the disease(1). Except for the Acute Promyelocytic Leukemia (APL) subtype, the standard treatment of AMLs has little changed over the past 40 years. It generally consists of intensive chemotherapy composed of one anthracyclin (daunorubicin or idarubicin) and the nucleoside analog cytarabine (Ara-C). However, relapses are frequent (40 to 70% of patients, depending on prognosis factors)(2) and the overall survival very low, making novel treatments urgently needed.

Differentiation therapies have appeared as a powerful strategy for AML treatment. They rely on the idea that restoration of differentiation is associated with cell division arrest, followed by death due to the naturally limited lifespan of differentiated cells. This approach has proved particularly efficient at curing APL(3), a minor AML subtype characterized by the expression of oncogenic fusion proteins engaging the retinoic acid receptor a (RARa), which belongs to the nuclear receptor family. APL therapy is based on pharmacological doses of its natural ligand, all*-trans*-retinoic acid (ATRA) used in combination with Arsenic Trioxide(4). ATRA leads to the degradation of the oncogenic RARα fusion protein and activates wild-type RARα. This activates a specific transcriptional program, which drives differentiation, cell cycle arrest and death of the leukemic cells(5,6). ATRA also induces to various degrees the *in vitro* differentiation of certain non-APL AML cell lines and primary cells from a substantial number of non-APL AMLs(6). This includes AMLs mutated in the NPM1-(7), IDH1/IDH2-(8) or FLT3-ITD genes(9) or overexpressing the transcription factor EVI-1(10). However, the clinical trials conducted so far have failed to prove a significant efficacy of ATRA on non-APL AML patients, including in combination with other drugs(11). This lack of effect has been attributed to the inability of ATRA to induce the expression of critical genes involved in differentiation, cell cycle arrest or apoptosis(6,12). Interestingly, targeting epigenetic enzymes has recently appeared promising to restore the ability of ATRA to activate RARα target genes and potentiate ATRA-induced differentiation of non-APL AMLs(6,11,12). For instance, inhibition of histone deacetylases (HDAC) with valproic acid was shown to favor ATRA-induced differentiation of non-APL AML cells(13). However, the clinical trials conducted so far showed only limited effects in a subset of the patients treated with the combination of ATRA and valproic acid(14–16). More recently, inhibition of the histone demethylase LSD1/KDM1A was also shown to strongly sensitize AML cells to ATRA in preclinical models *via* transcriptional reprogramming(17). Phase I/II trials are ongoing to determine the clinical efficacy of the combination between ATRA and LSD1 inhibitors.

SUMO is a group of 3 (SUMO-1 to -3) ubiquitin-related polypeptidic post-translational modifiers covalently and reversibly conjugated to numerous intracellular proteins to regulate their function and fate(18). Its conjugation involves a unique E1 SUMO-activating enzyme, a unique E2-conjugating enzyme (Ubc9) and several E3 SUMO ligases(19). Deconjugation is ensured by deSUMOylases, in particular of the SENP family. Increasing evidence links deregulation of the SUMO pathway to cancer(20,21), including in hematological malignancies such as lymphomas(22) and multiple myeloma(23). In the case of AMLs, the SUMO pathway is essential for efficient differentiation therapy of APLs by an ATRA + arsenic trioxide combination treatment. Arsenic trioxide induces the rapid SUMOylation of the PML-RARα oncoprotein, which initiates its elimination by the ubiquitin/proteasome system(24–26). In addition, we have shown that inhibition of the SUMO pathway by genotoxics-induced reactive oxygen species (ROS) is essential for fast and efficient cell death of chemosensitive non-APL AML cells subjected to antracyclins or Ara-C treatment(27).

SUMO is increasingly viewed as an epigenetic mark highly enriched at gene promoters(28–30). It regulates gene expression via modification of numerous transcription factors/co-regulators, histone-modifying enzymes, RNA polymerases and even histones(31,32). Although sometimes associated with transcriptional activation(33), SUMOylation at gene promoters is mostly known to limit or repress transcription(28,32,34–37). In particular SUMOylation facilitates the recruitment of SUMO-interacting motifs (SIM)-containing co-repressors on promoters(31,38,39). We show here that SUMOylation takes part in the epigenetic silencing of ATRA-responsive genes in non-APL AMLs. Its inhibition strongly activates the pro-differentiating and anti-leukemic effect of ATRA in these cancers, which opens new perspectives in the treatment of this poor prognosis cancer.

## Methods

### Cell lines and primary AML patient cell culture

U937, HL60, THP1, MOLM14 cells (obtained from DSMZ, Germany) were cultured in RPMI containing 10% fetal bovine serum (FBS) and streptomycin/penicillin at 37°C in the presence of 5% CO_2_. U937 cells resistant to Ara-C were generated by culturing the cells for 2 months in the presence of increasing Ara-C concentration (up to 0.1 *μ*M). Fresh bone marrow aspirates were collected after obtaining informed consents from patients (Ethical Committee « Sud Méditerranée 1 », ref 2013-A00260-45, HemoDiag collection). Fresh leukocytes were purified using density-based centrifugation using Histopaque 1077 from SIGMA and resuspended at a concentration of 10^6^/ml in IMDM (SIGMA) complemented with 1.5 mg/ml bovine serum albumin, 4.4 /*μ*g/ml insulin (SIGMA), 60 /*μ*g/ml transferrin (SIGMA), 5% streptomycin + penicillin, 5% FBS, 5 ß-mercaptoethanol, 1 mM pyruvate, 1x MEM non-essential amino acids (Life Technologies), 10 ng/ml IL-3 (PeproTech), 40 ng/ml SCF (PeproTech), and 10 ng/ml TPO (PeproTech).

### Pharmacological inhibitors, reagents, and antibodies

All-*trans*-retinoic acid (ATRA) was from Sigma. It was resuspended at a 100 mM concentration in DMSO and stored at −20°C for a maximum of 2 weeks. Anacardic acid was from Santa Cruz Biotechnologies and 2-D08 from Merck-Millipore. Anti-SUMO-1-(21C7) and SUMO-2 (8A2) hybridomas were from the Developmental Studies Hybridoma Bank. The anti H3K4Me_3_ antiserum was from Abcam.

### Flow cytometry

Cells were washed in PBS containing 2% FBS and incubated at 4°C for 30 minutes in the presence of the following fluorophore-conjugated antibodies: CD45-Pacific Blue (A74763; Beckman Coulter), CD14-PE (130-091-242; Miltenyi), CD15-PE-Vio770 (130-100-425; Miltenyi), CD11b-APC (130-109-286; Miltenyi). Matched isotype controls were used for each treatment condition. After washing, cells were analyzed using the LSR Fortessa flow cytometer (Becton Dickinson) and the FacsDiva software. Data were analyzed using the FlowJo software (version 10). For patient samples, MFIs for each differentiation marker were measured on leukemic cells previously selected using CD45/SSC gating(40). MFIs from isotype controls were subtracted from each treatment condition.

### Microscopic analyses

Cell lines or patient samples were cytospun on microscope slides (1500 rpm for 5 min), dried for 5 min and stained, first with May-Grunwald-(5 min) and, then, Giemsa (1/10 dilution, 15 min) stain (MGG staining). Microscopic examinations were performed using the AxioImager Z2 microscope (Zeiss).

### Cell viability, cell cycle and proliferation assay

For proliferation assays on cell lines, cells were seeded at a concentration of 3 × 10^5^/ml and viable cells were counted at regular intervals using the Trypan-blue exclusion method with an EVE automatic cell counter. Cell cycle distribution was analyzed by Propidium Iodine (PI) staining. Cells were washed once with PBS and fixed with cold 70% ethanol for 10 min and washed once with PBS. 100/*μ*g/mL RNAase A (Sigma) was then added for 10 min at room temperature. Cells were washed with PBS and stained with 50 /*μ*g/mL PI (Sigma) for 10 min at room temperature. Cells were then washed once with PBS and analyzed by flow cytometry. For patient cells, equal numbers of CountBright^TM^ absolute counting beads (C36950; Life Technologies) were added to each sample. Viable cells were selected using the CD45/SSC gating and their number was normalized to the number of beads counted in the same sample.

### Retroviral Infections

Retroviral constructs expressing either Ubc9, SENP2 or SENP5 were constructed by inserting human cDNA using the Gateway cloning technology (ThermoFisher Scientific) into the pMIG retroviral vector(41), which also coexpress EGFP from the same polycistronic mRNA. Retroviruses were produced by cotransfection of these constructs with gag-pol- and VSVG expression vectors into HEK293T cells using Lipofectamine-2000 (Invitrogen). Viral supernatants were collected 48hr later, 0.45 *μ*m-filtered and directly used to infect AML cell lines. Only EGFP-positive cells were considered in flow cytometry analyses. Where indicated, the EGFP-positive cells were sorted using the FACS-Aria cell sorter (Beckton Dickinson)

### RT-qPCR assays

Total mRNA was purified using the GenElute Mammalian Total RNA kit (Sigma). After DNase I treatment, 1 *μ*g of total RNA was used for cDNA synthesis using the Maxima First Strand cDNA kit (ThermoFisher Scientific). qPCR assays were conducted using Taq platinium (Invitrogen) and the LightCycler 480 device (Roche) with specific DNA primers (see primer table). Data were normalized to the housekeeping *TBP* mRNA levels.

### Chromatin immunoprecipitation assays (ChIPs)

30.10^6^ cells were cross-linked with 1% paraformaldehyde for 8 minutes. Paraformaldehyde was then neutralized with 125 mM glycine for 10 minutes. Cross-linked cells were washed with cold PBS, resuspended in a cell lysis buffer (5 mM PIPES pH7.5, 85 mM KCl, 0.5% NP40, 20 mM N-ethyl maleimide, 1 *μ*g/mL of aprotinin, pepstatin, leupeptin, 1 mM AEBSF) and incubated at 4°C for 10 minutes with rotation. Nuclei were centrifuged (5,000 rpm for 10 minutes at 4°C) and resuspended in a nucleus lysis buffer (50 mM Tris-HCl pH 7.5, 1% SDS, 10 mM EDTA, 20 mM N-ethyl maleimide, 1 *μ*g/mL of aprotinin, pepstatin, leupeptin, 1 mM AEBSF) and incubated at 4°C for 2.5 hours. Lysates were then sonicated for 30 cycles of 30 seconds each at 4°C using the Bioruptor Pico (Diagenode) under standard conditions. After sonication, samples were centrifuged (13,000 rpm for 10 minutes at 4°C) and the supernatants were diluted 10-fold in the immunoprecipitation buffer (1.1% Triton X100, 50 mM Tris-HCl pH 7.5, 167 mM NaCl, 5 mM N-ethyl maleimide, 1 mM EDTA, 0.01% SDS, 1 /*μ*g/mL of aprotinin, pepstatin, leupeptin, 1 mM AEBSF) with 2 *μ*g of antibodies and Dynabeads Protein G (ThermoFisher Scientific). Immunoprecipitations were performed overnight at 4°C. Beads were then washed in low salt buffer (50 mM Tris-HCl pH7.5, 150 mM NaCl, 1% Triton X100, 0.1% SDS, 1 mM EDTA), high salt buffer (50 mM Tris-HCl pH7.5, 500 mM NaCl, 1% Triton X100, 0.1% SDS, 1 mM EDTA), LiCl salt (20 mM Tris-HCl pH7.5, 250 mM LiCl, 1% NP40, 1% deoxycholic acid, 1 mM EDTA) and TE buffer (10 mM Tris-HCl pH7.5, 0.2% Tween20, 1 mM EDTA). Elution was done in 200 /*μ*L of 100 mM NaHCO_3_, 1% SDS. Chromatin crosslinking was reversed by overnight incubation at 65°C with 280 mM NaCl followed and 2h at 45°C with 35 mM Tris-HCl pH6.8, 9 mM EDTA, 88 *μ*g/mL RNAse, 88 *μ*g/mL Proteinase K. Immunoprecipitated DNAs were purified using the Nucleospin Gel and PCR Cleanup kit (Macherey-Nagel). Immunoprecipitated DNA and DNA inputs purified from samples before immunoprecipitation were subjected to PCR analysis using the Roche LightCycler 480 device using appropriate primers (see primer table)

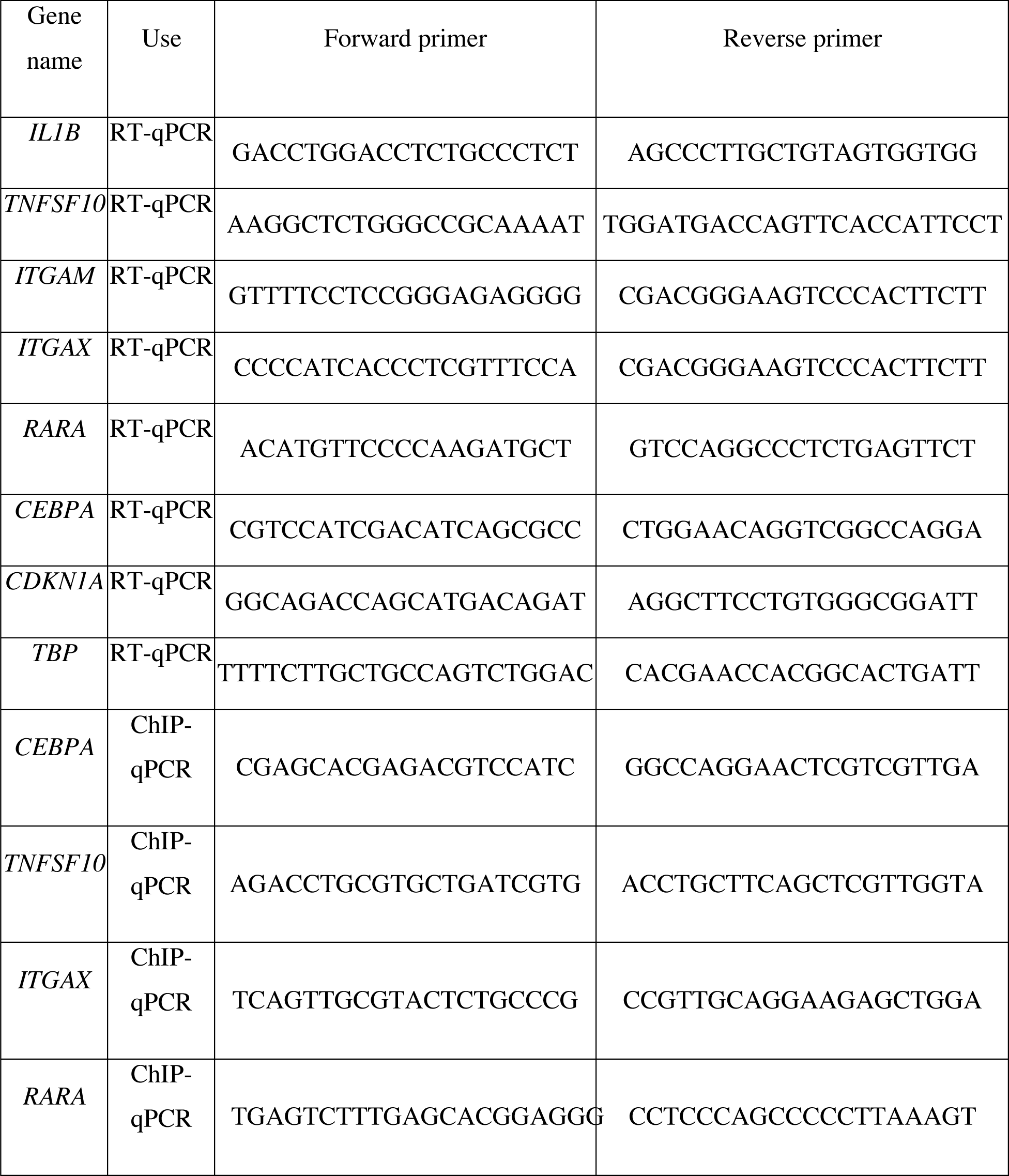

### Tumor xenografts

Animals were used in accordance to a protocol reviewed and approved by the Institutional Animal Care and Use Committee of Région Midi-Pyrénées (France). Tumors were generated by injecting subcutaneously 2 × 10^6^ U937 cells (contained in 100 *μ*l of PBS) on both flanks of NOD-Scid-IL2gRnull (NSG) mice (adult males and females; 25 g each; Charles River Laboratories). When tumors reached 100 mm^3^, mice received peritumoral injections of ATRA (2.5 mg/kg/day) or 2-D08 (10 mg/kg/day) or both every 2 days. Injections of vehicle (DMSO) were used as controls. Tumor sizes were measured with a caliper and volumes calculated using the formula: 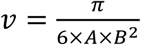, where A is the larger diameter and B is the smaller diameter.

### Statistical analysis

Statistical analyses were performed using the Prism 5 software. The two-tailed paired Student’s t test, the Wilcoxon matched-pairs signed rank test and the Mann-Whitney test were used for analysis of experiments with cell lines, patient samples and xenografts, respectively. Differences were considered as significant for p values of <0.05. *: p<0.05; **: p<0.01; ***: p<0.001. ****: p<0.0001. ns = non significant

## Results

### The SUMO pathway represses ATRA-induced differentiation of non-APL AML cells

To assess the role of SUMOylation in ATRA-induced differentiation of non-APL AMLs, we first resorted to the reference and well-characterized U937 and HL-60 non-APL AML cell lines. They belong to the M4 and M2 subtypes respectively (out of 8 AML subtypes of the French American British-FAB classification) and can differentiate *in vitro* in the presence of ATRA, albeit at low efficiency. We used two commercially available pharmacological inhibitors of the SUMO pathway: 2-D08, an inhibitor of the E2 SUMO-conjugating enzyme Ubc9(42,43) and anacardic acid (AA), an inhibitor of the SUMO-activating enzyme Uba2/Aos1(44). Both inhibitors were used at concentrations leading to moderate hypoSUMOylation of cell proteins (Figure 1A). This partial inhibition of the SUMO pathway resulted in a strong increase in ATRA-induced differentiation of both U937 (Figure 1B) and HL60 (Figure 1B) cells, as assessed by enhanced expression of CD15 or CD11b differentiation markers respectively. Combining ATRA with 2-D08 also increased the differentiation of THP1 (Figure 1D), MOLM14 (Figure 1E) and Ara-C-resistant U937 cells developed in our laboratory (Figure 1F). The combination of ATRA and SUMOylation inhibitors increased the number of U937 (Figure 2A), HL60 (Figure 2B) and THP1 (Supplementary Figure 1) cells showing morphological changes typical of terminal myeloid differentiation, such as nuclear lobulation and appearance of numerous cytosolic granules.

**Figure 1:**
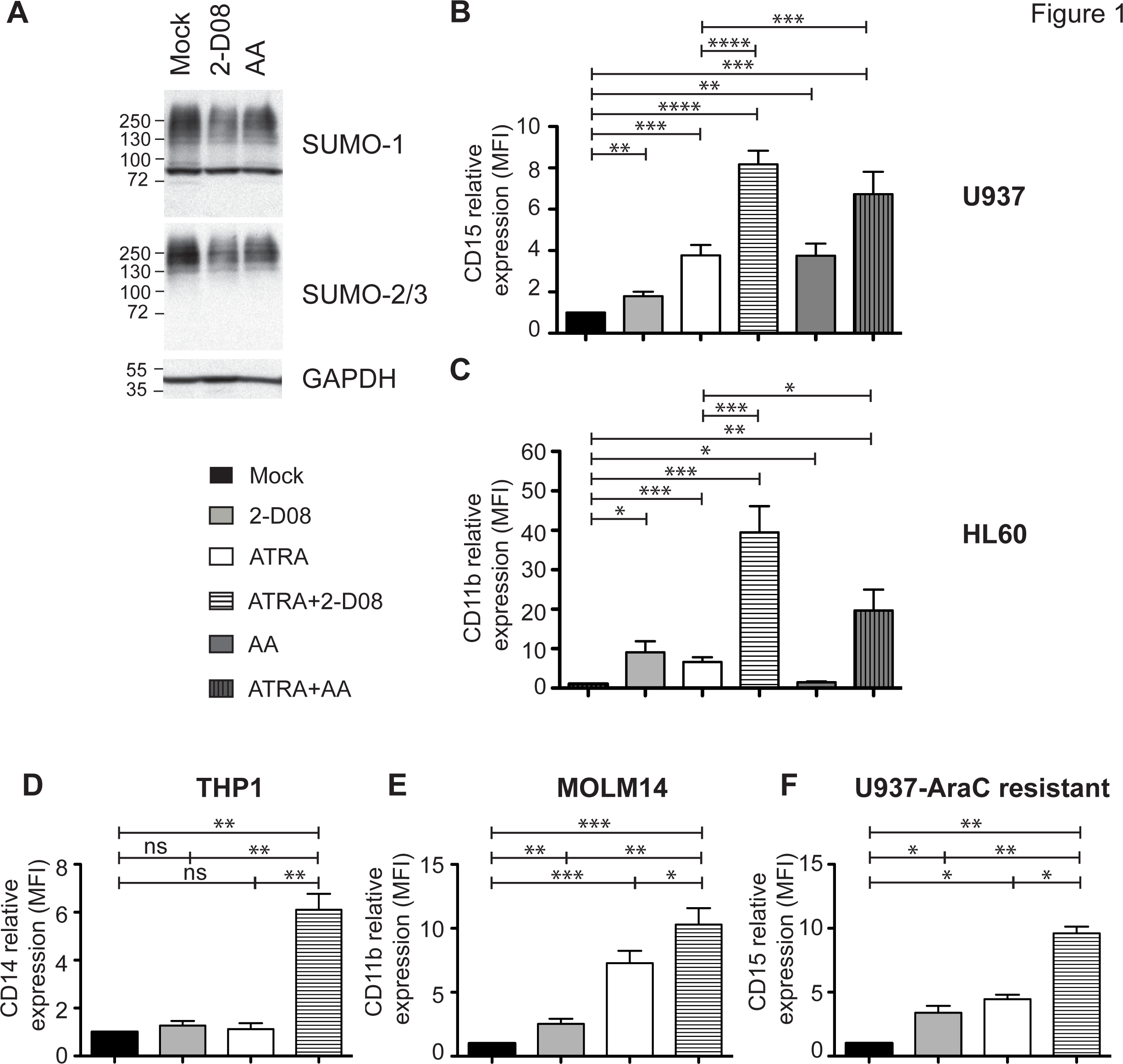
SUMOylation represses ATRA-induced differentiation of various non-APL AML cell lines, including chemoresistant ones. (A) *Reduction of cell protein SUMOylation in the presence of SUMO pathway inhibitors*. Cell extracts from U937 cells treated for 48 h with either 2-D08 (50 pM) or Anacardic Acid (AA) (25 pM) were used in immunoblotting experiments using antibodies directed to SUMO-1 and SUMO-2/3 (SUMO 2 and −3 being immunologically indistinguishable). SUMO-conjugated proteins appear as a smear, which is lighter in the presence of 2-D08 or AA. Immunoblotting against GAPDH was used as an electrophoresis loading control. (B) *Inhibitors of SUMOylation increase the differentiation of U937 cells*. U937 cells were treated for 5 days with ATRA (1 pM), 2-D08 (50 pM), AA (10 pM) or the combinations and the expression of CD15 was flow cytometry-assayed. Median fluorescent intensities (MFIs) normalized to mock-treated cells are indicated (mean +/−SEM n = 10). (C) *SUMOylation inhibitors increase the differentiation of HL60 cells*. HL60 cells were treated with ATRA (1 pM), 2-D08 (25 pM) or AA (25pM) alone or in combination for 5 days. CD11b expression was assayed by flow cytometry. MFIs, normalized to mock-treated cells are indicated on the graph (n = 11, mean +/−SEM) (D-F) *Differentiation of THP1, MOLM14 and Ara-C-resistant U937 cells*. Cells were treated for 9- (THP1), 2- (MOLM14) or 5 days (Ara-C-resistant U937) with ATRA (0.5 pM, 0.1 pM and 1 pM respectively), 2-D08 (50 pM) or ATRA+2-D08 and CD14, CD11b or CD15 expressions were assayed. MFIs, normalized to mock-treated cells are indicated on the graph (n = 4 for THP1, n = 7 for MOLM14, n = 3 for Ara-C resistant U937, mean +/−SEM).

**Figure 2:**
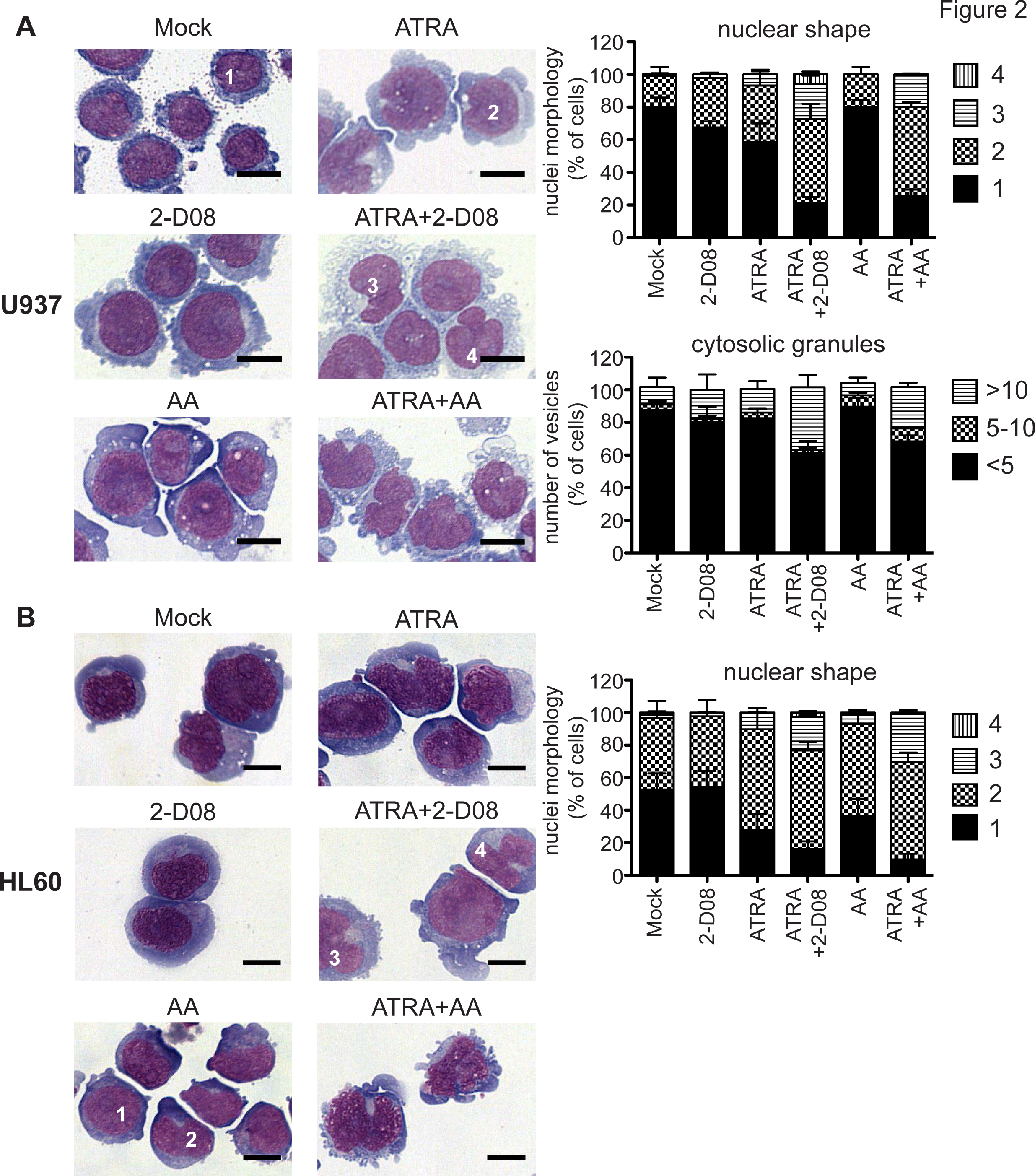
Inhibition of SUMOylation increases morphological changes associated with ATRA-induced differentiation. (A) *Morphological changes of U937 cells*. U937 cells were treated for 5 days with ATRA (1 pM), 2-D08 (50 pM), AA (10 pM) or the combinations and May-Grumwald-Giemsa (MGG)-stained before microscopic analysis. Scale bar = 10 pm (n = 3). Cells were distributed in 4 groups, depending on their nuclear morphology and 3 groups for the number of cytosolic granules. Quantification was done on at least 100 cells from 3 independent experiments. (B) *Morphological changes of HL-60 cells*. HL-60 cells were treated for 9 days with ATRA (1 pM), 2-D08 (25 pM) or AA (25pM) alone or in combination. Scale bar = 10 pm (n=3). Cells were distributed in 4 groups, depending on their nuclear morphology as in A. Quantification was done on at least 100 cells from 3 independent experiments.

Altogether, these data suggest that the SUMO pathway limits the differentiating effects of ATRA on various non-APL AML cell lines and that its inhibition could favor their ATRA-induced differentiation, including in the case of resistance to a chemotherapeutics used in AML treatment.

### SUMOylation limits ATRA-induced expression of myeloid differentiation-associated genes

We then wondered if the repressive action of SUMOylation on ATRA-induced differentiation could be linked to its well-characterized ability to repress/limit gene expression(38). Inhibition of SUMOylation with 2-D08 was sufficient on its own to increase the expression of various ATRA-responsive genes associated with myeloid differentiation, such as *RARA, CEBPA, TNFSF10, ITGAX, ITGAM* and *IL1B* (Figure 3A). This suggests that inhibition of SUMOylation might prime differentiation by increasing the basal expression of master genes involved in myeloid differentiation. In addition to basal expression, 2-D08 also increased the ATRA-induced expression of most of those genes (Figure 3A). To determine if SUMOylation controls the expression of these genes at the level of chromatin, we assayed the active transcription-associated histone mark H3K4me_3_ on their promoters. An increase of this mark correlated with the level of expression of *RARA, 1TGAX, CEBPA* and *TNFSF10* mRNAs (Figure 3B), suggesting that SUMOylation represses ATRA-responsive genes induction at the level of chromatin.

**Figure 3:**
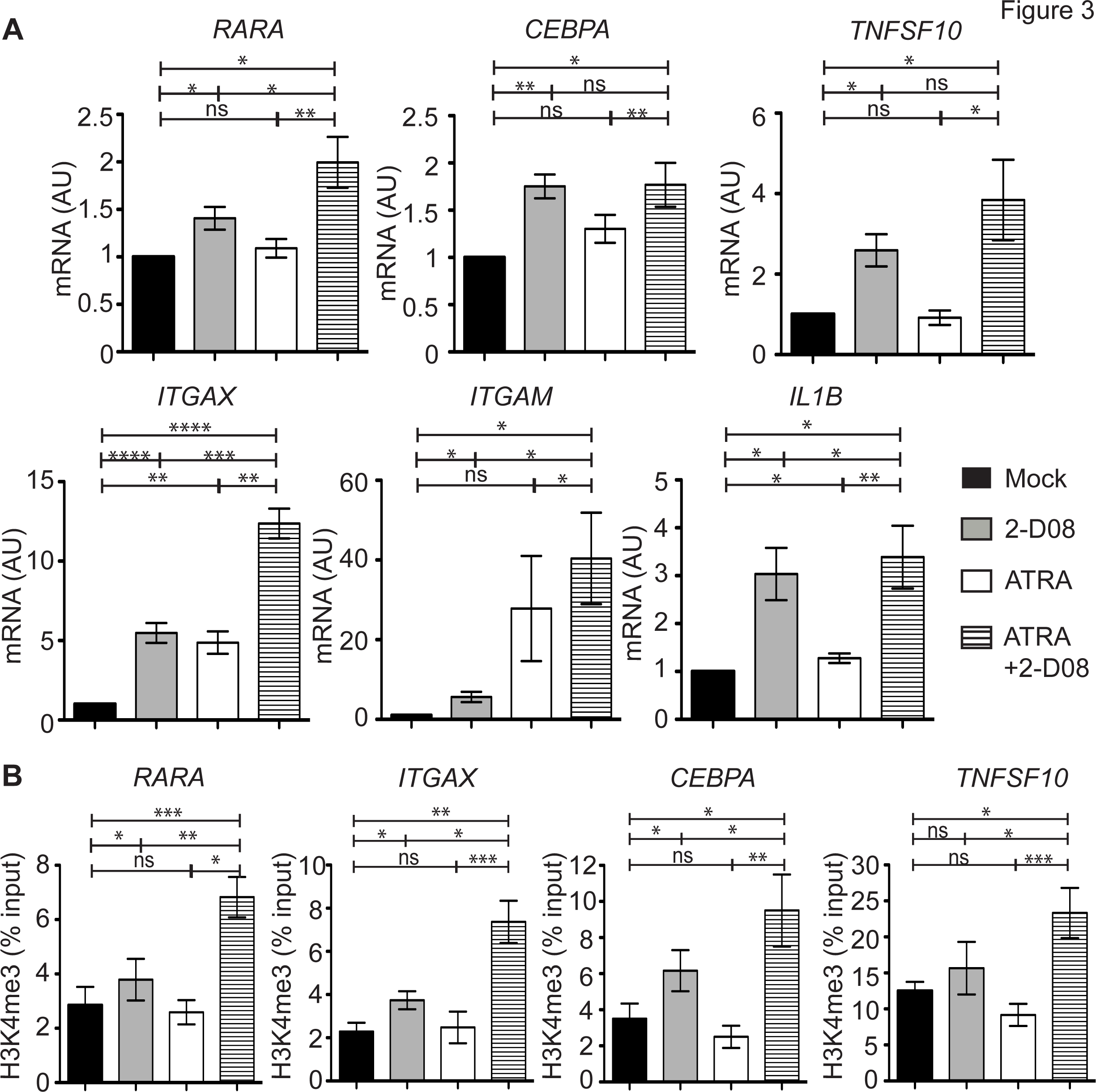
SUMOylation silences ATRA-responsive genes at the level of chromatin. (A) *Expression of differentiation-associated genes*. U937 cells were treated for 2 days with ATRA (1 μM), 2-D08 (50 μM) or ATRA+2-D08. mRNAs for the indicated genes were assayed by qRT-PCR and normalized to the TBP housekeeping mRNA. The results are presented as ratios to mock-treated cells (n = 6, mean +/− SEM). (B) *Chromatin immunoprecipitation assays*. U937 cells were treated as in (A) and used in ChIP experiments using control- and H3K4Me_3_ antibodies. Genomic regions showing the highest enrichment of H3K4me3 in the ENCODE database on the *RARA, ITGAX, CEBPA and TNFSF10* promoters in PBMCs were amplified from the immunoprecipitated DNA fragments (see primer list). The results are presented as percentage of input. The signal from the control IP was subtracted (n = 5, mean +/− SEM).

### Inhibition of SUMOylation potentiates the anti-leukemic effects of ATRA

Differentiated cells stop proliferating and have a shorter lifespan than undifferentiated cells. Consistently, ATRA and 2-D08 synergized to block the proliferation of both U937 cells (Figure 4A) and their Ara-C resistant variants *in vitro* (Figure 4B). This correlated with an accumulation of cells in G0/G1 (Figure 4C) and a strong activation of the *CDKN1A* gene encoding the CDK inhibitor p21CIP1 (Figure 4D). This suggests that inhibition of SUMOylation increases the anti-proliferative effects of ATRA *in vitro* by inducing a cell cycle arrest. To determine if this would also be the case *in vivo*, we treated immunodeficient mice subcutaneously xenografted with U937 cells with ATRA, 2-D08 or ATRA+2-D08 after engraftment. Only the combination of ATRA and 2-D08 induced a significant reduction in tumor growth whereas ATRA and 2-D08 alone showed slight, if any, effects (Figure 4E-F). Thus, the combination of ATRA with an inhibitor of SUMOylation can, not only promote non-APL AML cell differentiation, but also exert anti-proliferative effects both *in vitro* and *in vivo*.

**Figure 4:**
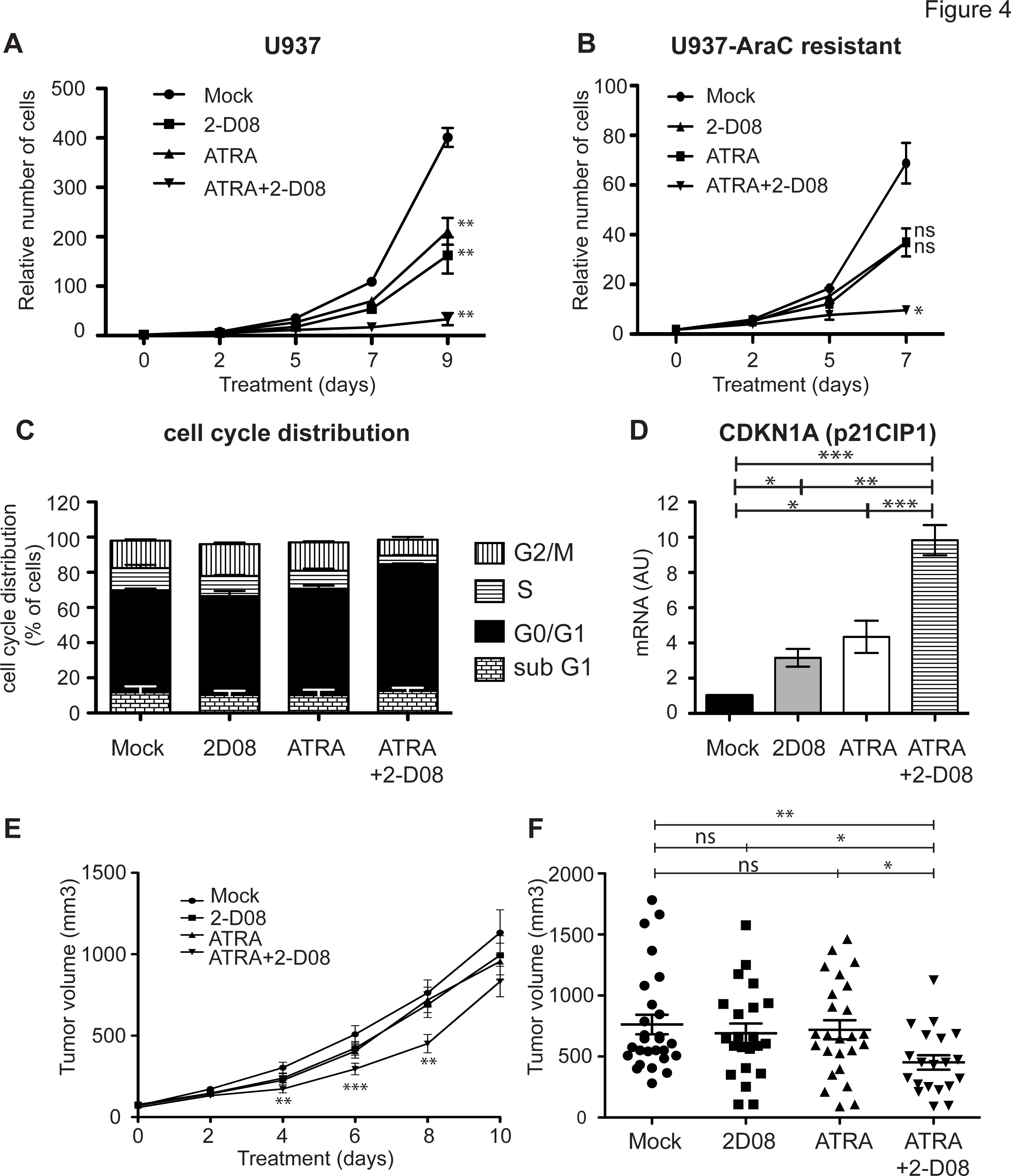
Inhibition of SUMOylation with 2-D08 reduces AML cells proliferation *in vitro* and *in vivo*. (A-B) *Cell proliferation*. Parental or Ara-C-resistant U937 cells were treated with ATRA (1 pM), 2-D08 (50 pM) or the combinations for the indicated times. Viable cells were quantified using Trypan blue exclusion. The number of cells on Day 0 was set to 1. The p-values are between the mock-treated and treated cells (n = 3, mean +/− SEM). (C) *Cell cycle distribution*. U937 cells were treated as in A for 9 days and cell cycle distribution was monitored using propidium iodine (PI) staining (n=4, mean +/− SEM). (D) *Increased expression of p21CIP1*. U937 cells were treated with ATRA (1 pM), 2-D08 (50 pM) or ATRA+2-D08 for 2 days. mRNAs for *CDKN1A* was assayed by qRT-PCR and normalized to the TBP housekeeping mRNA. The results are presented as ratios to mock-treated cells (n = 6, mean +/− SEM). (E-F) *Xenografts growth*. U937 cells were xenografted subcutaneously on each flank of NSG mice (2 independent experiments, 7 mice per group in each experiment). Four days after injection, mice were treated peritumorally with 2-D08 (10 mg/kg) and/or ATRA (2.5 mg/kg) every 2 days for 10 days. Tumor volumes were measured every 2 days and are presented for all mice at day 8 on (F). Tumors smaller than 30 mm^3^ before the beginning of the treatment were excluded from the analysis. The p values indicated in (E) were calculated between the DMSO and the ATRA+2-D08 conditions.

### Genetic modulation of the SUMO pathway affects ATRA-induced differentiation of non-APL AML cells

To rule out possible off-target effects of 2-D08 and anacardic acid, we resorted to the genetic manipulation of the SUMO pathway to confirm the role of SUMOylation on ATRA-induced differentiation in non-APL AMLs. In a first step, we induced a hypo-SUMOylated state in U937 cells by overexpressing either the SENP-2- or the SENP-5 desumoylase. This led to a significant increase in their ATRA-induced differentiation, as assayed by the expression of CD11b (Figure 5A). Interestingly, this correlated with a reduction in their proliferation (Figure 5B). Overexpression of SENP-2 also increased ATRA-induced differentiation of HL-60 cells (Figure 5C) and was associated with stronger expression of various ATRA-responsive genes *(RARA, CEBPA, 1TGAM* and *1L1B;* Figure 5D). Interestingly, although SENP-2-expressing HL60 cells did not appear more differentiated than control cells in the absence of ATRA, they showed higher basal expression of these genes. This further supports the idea that inhibition of SUMOylation could prime AML cells for differentiation. In a second step, we increased SUMOylation in THP1 cells via overexpression of the SUMO E2-conjugating enzyme Ubc9. This led to a massive decrease in ATRA-induced differentiation, as assayed by CD14 expression (Figure 6A) and morphological changes (appearance of pseudopods, cytosolic vesicles and granules) (Figure 6B). Moreover, Ubc9 overexpression strongly decreased basal expression and/or induction of genes involved in myeloid differentiation by ATRA (Figure 6C). Altogether, this confirms the repressive role of the SUMO pathway on ATRA-induced differentiation of non-APL AMLs.

**Figure 5:**
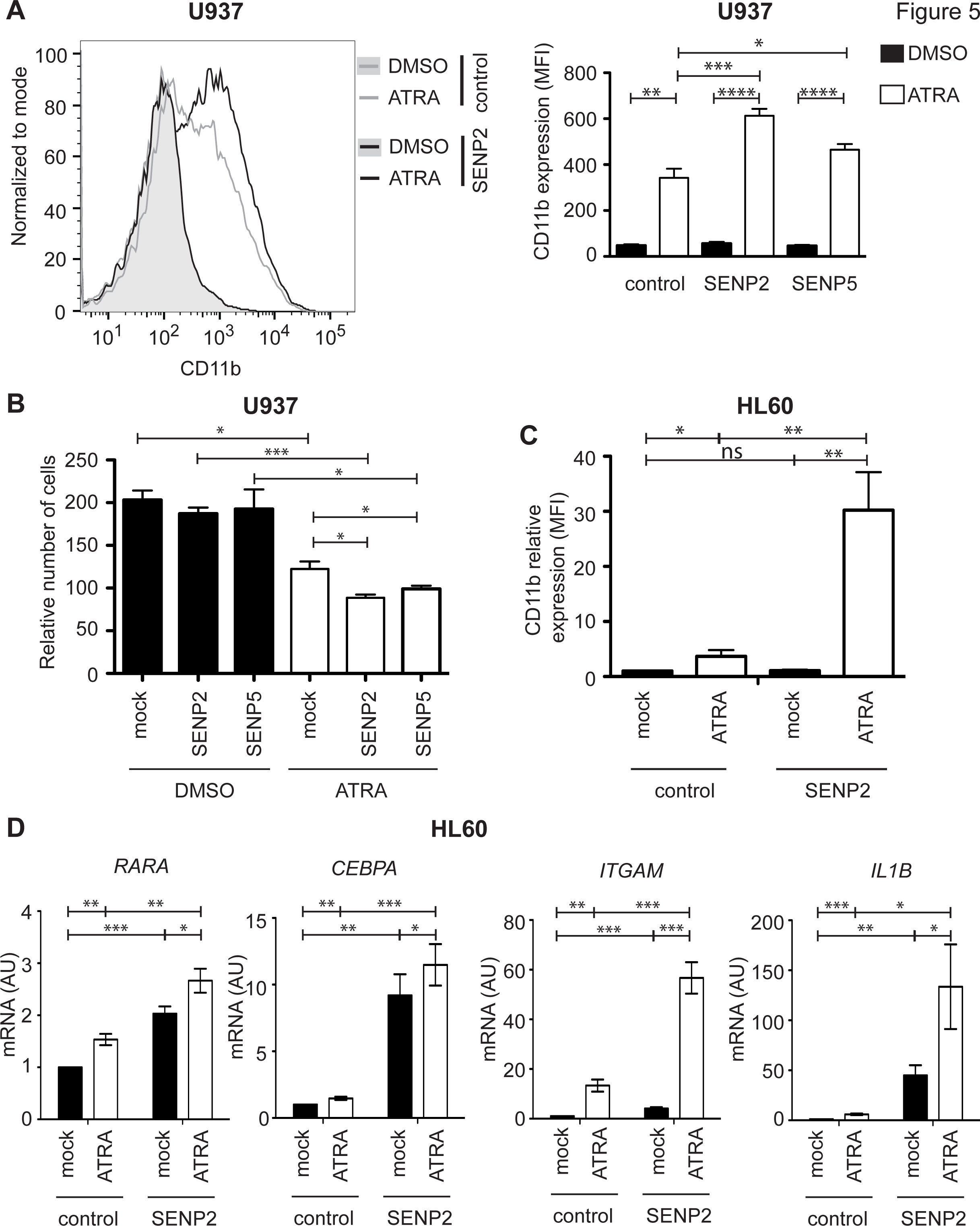
Overexpression of SENP desumoylases favors ATRA-induced differentiation and anti-leukemic activity. (A) *Differentiation of U937 cells*. U937 cells were infected with either a control- or a SENP2- or SENP5-expressing retroviruses also expressing EGFP. EGFP-expressing cells were FACS-sorted and treated, or not, with ATRA (1 μM) for 5 days, and assayed for the expression of CD11b. A representative histogram and averaged CD11b MFIs are presented on the left and right panels, respectively (n = 3, mean +/− SEM). (B) *U937 cell proliferation*. The same cells as in (A) were treated with ATRA (1 μM) for 9 days, at which time viable cells were scored. The number of cells on Day 0 was set to 1 (n = 4, mean +/− SEM). (C) *Differentiation of HL-60 cells*. HL-60 cells infected with a control- or a SENP2-expressing retrovirus and sorted for GFP expression were treated with ATRA (1 μM) for 2 days. CD11b expression (MFI) normalized to mock-treated control cells is shown (mean +/− SEM, n = 11). (D) *Expression of differentiation-associated genes in HL-60 cells*. mRNAs were purified from the cells used in C and assayed by qRT-PCR for the indicated genes and normalized to TBP mRNA. The results are presented as ratios to control conditions (DMSO-treated cells infected with the control retrovirus) (n = 6, mean +/− SEM).

**Figure 6:**
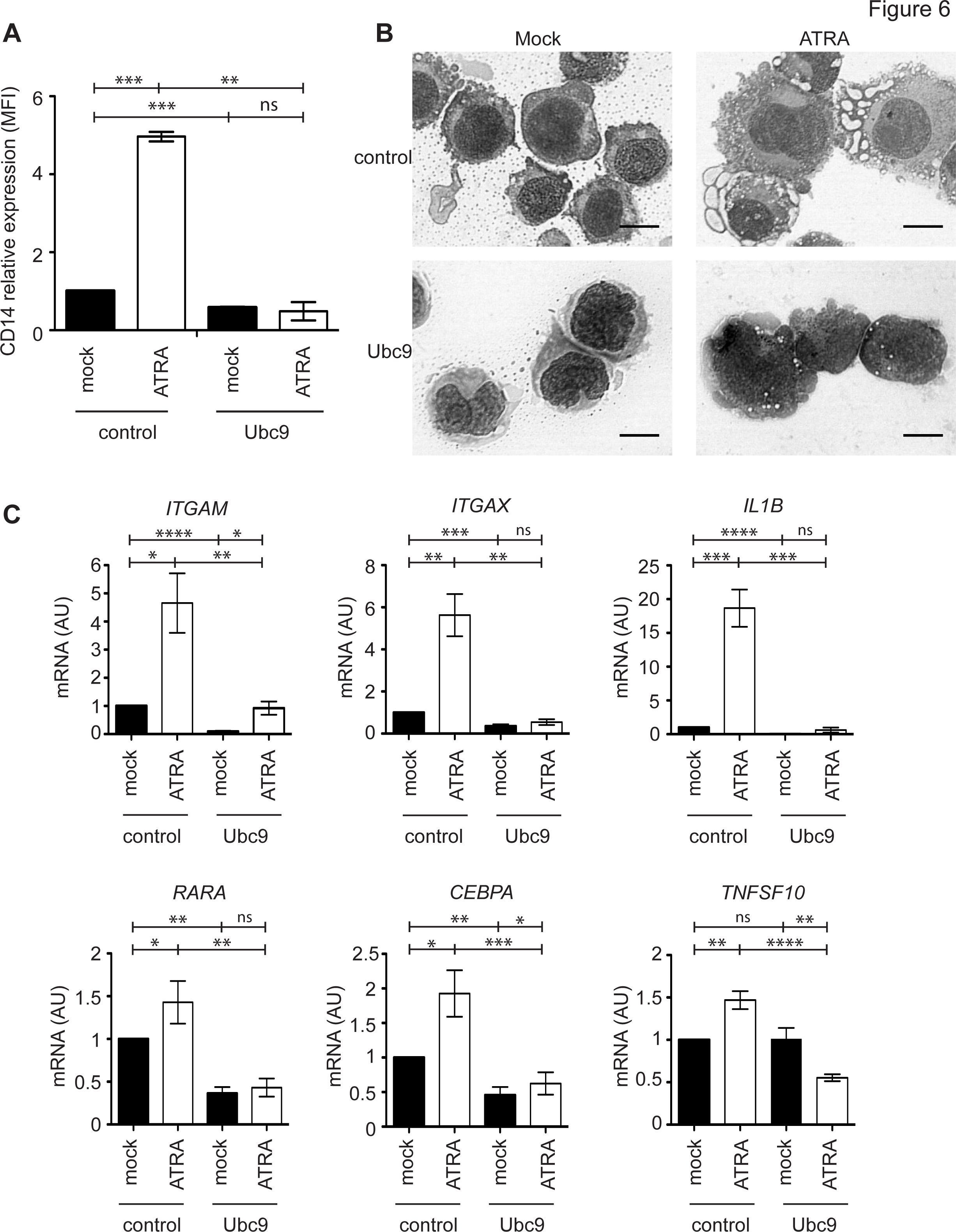
Overexpression of the E2 SUMO-conjugating enzyme limits ATRA-induced differentiation of THP1 cells. THP1 cells were infected with control- or Ubc9-expressing retroviruses also expressing EGFP for FACS sorting. (A) *THP1 cell differentiation*. Transduced cells were treated, or not, with ATRA (1 μM) for 9 days. Note that the ATRA concentration was twice that used in Figure 2A to induce differentiation. Differentiation was assessed by flow cytometry assay of CD14 expression and normalized to the mock-treated control cells (n = 3, mean +/− SEM). (B) *THP1 cell morphological changes*. Transduced cells were treated or not with ATRA (1 μM) for 5 days and their morphological differentiation was assessed after MGG staining. Scale bar = 10 μm. (C) *Expression of differentiation-associated genes*. Transduced cells were treated with ATRA (1 μM) for 5 days and the expression of mRNAs for the indicated genes was qRT-PCR-assayed and normalized to the TBP mRNA (n = 7, mean +/− SEM).

### Inhibition of SUMOylation potentiates the pro-differentiating and anti-proliferative activities of ATRA on primary AML cells

We finally asked whether inhibiting SUMOylation could also favor the *in vitro* differentiation of primary AML cells using bone marrow aspirates from AML patients at diagnosis. ATRA alone did not significantly induce their differentiation, as assayed by CD15 expression. 2-D08 and AA alone showed a slight pro-differentiating trend with, however, no statistical significance. In contrast, combining either of them with ATRA significantly increased CD15 expression compared to cells treated with ATRA alone (Figure 7A). Some patient cells were more sensitive to the differentiating effects of the ATRA+SUMOylation inhibitor combination than others, but they neither belonged to a unique FAB subtype nor shared cytogenetic/genetic abnormalities tested at diagnosis. Inhibitors of SUMOylation also increased the number of cells showing morphological changes typical of differentiation, such as nuclear lobulation, cytosol enlargement or appearance of cytosolic granules on cells from the two patients tested (Figure 7B and Supplementary Figure 2). Interestingly, 2-D08 and AA also potentiated ATRA-induced differentiation of primary cells from 1 patient not responsive to induction chemotherapy (AML1), as well as from 2 (AML2 and −4) out of 3 patients at relapse (Figure 7C). Importantly, both inhibitors increased the anti-leukemic activity of ATRA on the primary AML cells (Figure 7D). Altogether, these data confirm that SUMOylation inhibitors potentiate ATRA-induced differentiation of different non-APL AML subtypes and indicated novel therapeutic approach, including in the case of conventional chemotherapy failure.

**Figure 7:**
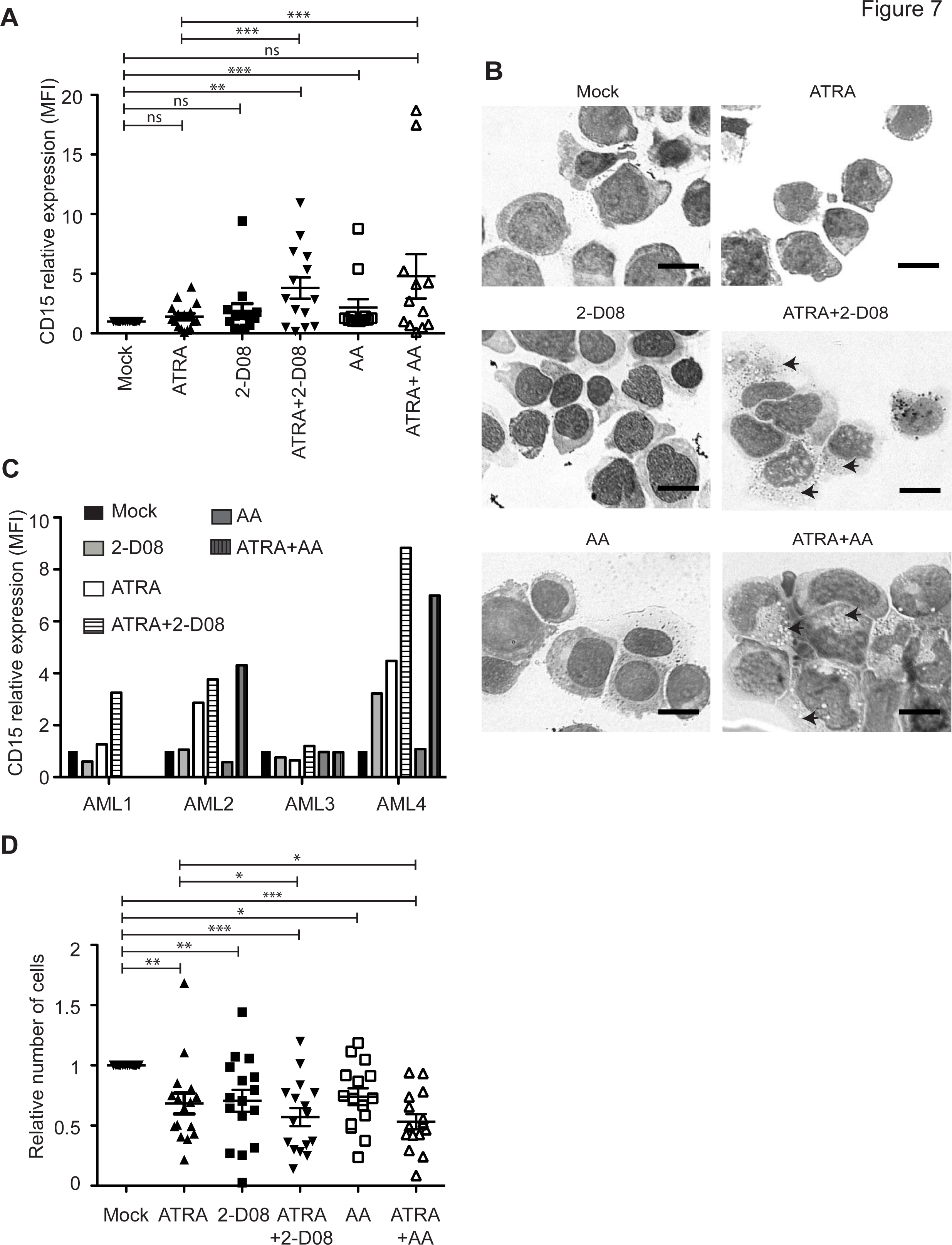
Inhibition of the SUMO pathway induces both differentiation and death of primary non-APL AML cells. (A) *CD15 expression on primary AML samples*. Primary AML cells from 16 patients were purified from bone marrow aspirate at diagnosis and treated *in vitro* with ATRA (1 μM), 2-D08 (50 μM), AA (25μM) or combinations of these drugs. Their differentiation was assessed 9 days later by flow cytometry assay of CD15. MFIs for each condition were corrected with isotypic controls and normalized to the mock condition (DMSO-treated samples). (B) *Morphological changes*. The bone marrow aspirate from one AML patient was treated as in (A) for 9 days and subjected to microscopic analysis after MGG staining. Scale bar = 10 μm. The arrows indicate cytosolic granules (also see Supplementary Figure 3) (C) *Analysis of non-responsive or relapsed patient samples*. Primary cells from 4 patients were treated and analyzed as in (A). AML-1 corresponds to a patient refractory to induction chemotherapy and AML-2 to −4 to patients at relapse. (D) *Viability of patient cells*. Primary AML cells taken at diagnosis or relapse (16 patients) were treated *in vitro* with ATRA (0.1 μM), 2-D08 (50 μM), AA (25 μM) or combinations of these drugs. The relative numbers of viable leukemic cells (CD45/SSC gating) were quantified 9 days later by flow cytometry using counting beads.

## Discussion

Differentiation therapies using ATRA have transformed APL from a fatal to a highly curable disease. Unfortunately, their efficacy in other AML subtypes has been disappointing. Our work demonstrates that SUMOylation represses ATRA-induced differentiation of non-APL AMLs. Mechanistically, the SUMO pathway silences genes critical for myeloid differentiation *(RARA* and *CEBPA)*, as well as genes required for cell cycle arrest *(CDKN1A)* and retinoid-induced apoptosis *(IL1B* and *TNFSF10)*. Inhibition of SUMOylation increases the presence of H3K4me3, a mark of active transcription, on their promoters and enhances both their basal and ATRA-induced expression. This suggests that targeting SUMOylation could prime AML cells for ATRA-induced differentiation, notably by augmenting the expression of critical regulators of myeloid differentiation.

More than 6000 SUMOylated proteins have been identified so far(18,45). Among them are many transcription factors and co-regulators, some of which play key roles in myeloid differentiation. This is the case of CEBPa and CEBPe. Their SUMOylation was shown to repress and activate their transactivation capacities, respectively(46,47). The ATRA-receptor RARα also undergoes dynamic SUMOylation/deSUMOylation cycles essential for its ATRA-induced activation(48,49). The SUMOylation of these transcription factors could participate in the silencing of ATRA-responsive genes that we have uncovered in non-APL AMLs. However, transcriptional repression of differentiation-associated genes by the SUMO pathway might not just result from the SUMOylation of single transcription factors, but from coordinated SUMOylation of multiple proteins bound to regulatory elements in the promoter/enhancers of these genes. Supporting the latter possibility, the concept of “group SUMOylation”, which has originally emerged from DNA repair regulation studies(50), implies that SUMO can exert its function wherever it is conjugated within a complex comprising several SUMOylatable proteins. SUMOylation of promoter-bound complexes are thought to favor, through SUMO/SIM interactions, the recruitment of transcriptional repressors(51) such as the N-Cor/HDAC(52) or the CoRest/LSD1(53) complexes. Inhibition of SUMO conjugation would lead to a decrease in their presence on the chromatin and would *in fine* facilitate both basal and ATRA-induced expression of genes involved in myeloid differentiation.

Our work indicates that combining ATRA with pharmacological inhibitors of the SUMO pathway strongly potentiates the pro-differentiating and anti-proliferative/pro-apoptotic effect of ATRA on non-APL AMLs including those resistant to the genotoxics currently used in the clinic. Considering the multiplicity of essential pathways controlled by SUMO, complete inhibition of SUMOylation would likely be too toxic in patients. However, similar to experiments with AA we conducted in a previous work(27), we observed no general toxicity of 2-D08 in mice. Both 2-D08 and AA are poorly efficient inhibitors of SUMOylation and induced only a slight decrease in SUMO conjugation at the doses we used. This probably explains their low toxicity. Finally, hemizigous mice expressing 50% of Ubc9 and showing slightly reduced SUMOylation activity are viable with no overt phenotype(54). This further suggests that limited reduction in cellular SUMOylation is not detrimental to essential cellular functions. Controlled inhibition of the SUMO pathway could thus, in combination with ATRA, target leukemic cells without overly affecting normal cells. In our experiments, the ATRA+2-D08 combination reduced AML growth *in vivo* but did not completely block it. The most likely explanation for this limited effect is that 2-D08, similarly to AA, has a poor bioavailability linked to its hydrophobic nature. Novel, more soluble SUMOylation inhibitors, or improvement of the pharmacological properties of the existing ones, are therefore necessary. ML-792 has been recently described as a new inhibitor of SUMOylation(55). However, its pharmacological properties and bioavailability have not been tested *in vivo*. Should they be better than those of 2-D08 and AA, this would permit an in-depth assessment of the therapeutic benefit of the ATRA+SUMOylation inhibitor association and, *in fine*, clinical use. In conclusion, our work suggests that targeting the SUMO pathway is a promising strategy to enhance the clinical efficacy of ATRA in non APL-AML and improve the treatment of this poor prognosis cancer.

## Acknowledgments

The authors declare no conflicting financial interests. We are grateful to all members of the “Oncogenesis and Immunotherapy” group of IGMM for support and fruitful discussions and to Pierre Gâtel, Rosa Paolillo and Frédérique Brockly for technical help. We thank Dr Robert Hipskind for critical reading of the manuscript. Funding was provided by the CNRS, Ligue Nationale contre le Cancer, FRM (Contract FDT20160435412), Cancéropole GSO, Région Languedoc-Roussillon (Contract Chercheur d’Avenir), INCA (ROSAML and METAML), Association Laurette Fugain (contract ALF-2017/02), the EpiGenMed Labex in the frame of the program “Investissement d’avenir” (ANR-10-LABX-12-01) and the Plan Cancer (to JK). The collection of clinical data and samples (HEMODIAG_2020) at the CHU Montpellier was supported by funding from the Région Languedoc-Roussillon and the SIRIC Montpellier Cancer.

## Authorship contributions

HB designed the experiments and performed most of the experimental work. MB performed some of the RT-qPCR assays and generated HL-60 cells expressing SENP2. JK performed the microscopic analyses and some of the flow cytometry experiments. TS generated THP1 cells lines expressing Ubc9. MH, SZ and JES performed the xenograft experiments. YH and GC collected and managed patient samples. MP and GB designed the study and wrote the manuscript.

## References

1. Döhner H, Weisdorf DJ, Bloomfield CD. Acute Myeloid Leukemia. N Engl J Med. 2015;373:1136–52

2. Dombret H, Gardin C. An update of current treatments for adult acute myeloid leukemia. Blood. 2016;127:53–61

3. Lo-Coco F, Avvisati G, Vignetti M, Thiede C, Orlando SM, Iacobelli S, et al. Retinoic acid and arsenic trioxide for acute promyelocytic leukemia. N Engl J Med. 2013;369:111–21

4. Ng C-H, Chng W-J. Recent advances in acute promyelocytic leukaemia. F1000Research [Internet]. 2017 [cited 2017 Sep 28];6. Available from: http://www.ncbi.nlm.nih.gov/pmc/articles/PMC5538034/

5. Gocek E, Marcinkowska E. Differentiation Therapy of Acute Myeloid Leukemia. Cancers. 2011;3:2402–20

6. van Gils N, Verhagen HJMP, Smit L. Reprogramming acute myeloid leukemia into sensitivity for retinoic-acid-driven differentiation. Exp Hematol [Internet]. 2017 [cited 2017 Jul 6]; Available from: http://www.sciencedirect.com/science/article/pii/S0301472X1730139X

7. Hajj HE, Dassouki Z, Berthier C, Raffoux E, Ades L, Legrand O, et al. Retinoic acid and arsenic trioxide trigger degradation of mutated NPM-1 resulting in apoptosis of AML cells. Blood. 2015;blood-2014-11-612416.

8. Boutzen H, Saland E, Larrue C, Toni F de, Gales L, Castelli FA, et al. Isocitrate dehydrogenase 1 mutations prime the all-trans retinoic acid myeloid differentiation pathway in acute myeloid leukemia. J Exp Med. 2016;213:483–97

9. Ma HS, Greenblatt SM, Shirley CM, Duffield AS, Bruner JK, Li L, et al. All-trans retinoic acid synergizes with FLT3 inhibition to eliminate FLT3/ITD+ leukemia stem cells in vitro and in vivo. Blood. 2016;127:2867–78

10. Verhagen HJMP, Smit MA, Rutten A, Denkers F, Poddighe PJ, Merle PA, et al. Primary acute myeloid leukemia cells with overexpression of EVI-1 are sensitive to all-*trans* retinoic acid. Blood. 2016;127:458–63

11. Ma HS, Robinson TM, Small D. Potential role for all- *trans* retinoic acid in nonpromyelocytic acute myeloid leukemia. Int J Hematol Oncol [Internet]. 2017 [cited 2017 Apr 15]; Available from: http://www.futuremedicine.com/doi/10.2217/ijh-2016-0015

12. Johnson DE, Redner RL. An ATRActive future for differentiation therapy in AML. Blood Rev. 2015;29:263–8

13. Trus MR, Yang L, Suarez Saiz F, Bordeleau L, Jurisica I, Minden MD. The histone deacetylase inhibitor valproic acid alters sensitivity towards all trans retinoic acid in acute myeloblastic leukemia cells. Leukemia. 2005;19:1161–8

14. Bug G, Ritter M, Wassmann B, Schoch C, Heinzel T, Schwarz K, et al. Clinical trial of valproic acid and all-trans retinoic acid in patients with poor-risk acute myeloid leukemia. Cancer. 2005;104:2717–25

15. Kuendgen A, Knipp S, Fox F, Strupp C, Hildebrandt B, Steidl C, et al. Results of a phase 2 study of valproic acid alone or in combination with all-trans retinoic acid in 75 patients with myelodysplastic syndrome and relapsed or refractory acute myeloid leukemia. Ann Hematol. 2005;84:61–6

16. Ryningen A, Stapnes C, Lassalle P, Corbascio M, Gjertsen B-T, Bruserud Ø. A subset of patients with high-risk acute myelogenous leukemia shows improved peripheral blood cell counts when treated with the combination of valproic acid, theophylline and all-trans retinoic acid. Leuk Res. 2009;33:779–87

17. Schenk T, Chen WC, Göllner S, Howell L, Jin L, Hebestreit K, et al. Inhibition of the LSD1 (KDM1A) demethylase reactivates the all-trans-retinoic acid differentiation pathway in acute myeloid leukemia. Nat Med. 2012;18:605–11

18. Hendriks IA, Vertegaal ACO. A comprehensive compilation of SUMO proteomics. Nat Rev Mol Cell Biol. 2016;17:581–95

19. Pichler A, Fatouros C, Lee H, Eisenhardt N. SUMO conjugation - a mechanistic view. Biomol Concepts. 2017;8:13–36

20. Seeler J-S, Dejean A. SUMO and the robustness of cancer. Nat Rev Cancer. 2017;

21. Eifler K, Vertegaal ACO. SUMOylation-Mediated Regulation of Cell Cycle Progression and Cancer. Trends Biochem Sci. 2015;40:779–93

22. Hoellein A, Fallahi M, Schoeffmann S, Steidle S, Schaub FX, Rudelius M, et al. Myc-induced SUMOylation is a therapeutic vulnerability for B-cell lymphoma. Blood. 2014;124:2081–90

23. Driscoll JJ, Pelluru D, Lefkimmiatis K, Fulciniti M, Prabhala RH, Greipp PR, et al. The sumoylation pathway is dysregulated in multiple myeloma and is associated with adverse patient outcome. Blood. 2010;115:2827–34

24. Tatham MH, Geoffroy M-C, Shen L, Plechanovova A, Hattersley N, Jaffray EG, et al. RNF4 is a poly-SUMO-specific E3 ubiquitin ligase required for arsenic-induced PML degradation. Nat Cell Biol. 2008;10:538–16

25. Lallemand-Breitenbach V, Jeanne M, Benhenda S, Nasr R, Lei M, Peres L, et al. Arsenic degrades PML or PML-RARα through a SUMO-triggered RNF4/ubiquitin-mediated pathway. Nat Cell Biol. 2008;10:547–55

26. Jeanne M, Lallemand-Breitenbach V, Ferhi O, Koken M, Le Bras M, Duffort S, et al. PML/RARA oxidation and arsenic binding initiate the antileukemia response of As2O3. Cancer Cell. 2010;18:88–98

27. Bossis G, Sarry J-E, Kifagi C, Ristic M, Saland E, Vergez F, et al. The ROS/SUMO Axis Contributes to the Response of Acute Myeloid Leukemia Cells to Chemotherapeutic Drugs. Cell Rep. 2014;7:1815–23

28. Neyret-Kahn H, Benhamed M, Ye T, Gras SL, Cossec J-C, Lapaquette P, et al. Sumoylation at chromatin governs coordinated repression of a transcriptional program essential for cell growth and proliferation. Genome Res. 2013;23:1563–79

29. Seifert A, Schofield P, Barton GJ, Hay RT. Proteotoxic stress reprograms the chromatin landscape of SUMO modification. Sci Signal. 2015;8:rs7–rs7.

30. Liu H, Zhang J, Heine GF, Arora M, Ozer HG, Onti-Srinivasan R, et al. Chromatin modification by SUMO-1 stimulates the promoters of translation machinery genes. Nucleic Acids Res [Internet]. 2012 [cited 2012 Oct 15]; Available from: http://nar.oxfordjournals.org.gate1.inist.fr/content/early/2012/08/30/nar.gks819

31. Raman N, Nayak A, Muller S. The SUMO system: a master organizer of nuclear protein assemblies. Chromosoma. 2013;122:475–85

32. Hendriks IA, Treffers LW, Verlaan-de Vries M, Olsen JV, Vertegaal ACO. SUMO-2 Orchestrates Chromatin Modifiers in Response to DNA Damage. Cell Rep. 2015;10:1778–91

33. Chymkowitch P, P A N, Aanes H, Robertson J, Klungland A, Enserink JM. TORC1-dependent sumoylation of Rpc82 promotes RNA polymerase III assembly and activity. Proc Natl Acad Sci. 2017;114:1039–44

34. Nayak A, Viale-Bouroncle S, Morsczeck C, Muller S. The SUMO-Specific Isopeptidase SENP3 Regulates MLL1/MLL2 Methyltransferase Complexes and Controls Osteogenic Differentiation. Mol Cell. 2014;55:47–58

35. Decque A, Joffre O, Magalhaes JG, Cossec J-C, Blecher-Gonen R, Lapaquette P, et al. Sumoylation coordinates the repression of inflammatory and anti-viral gene-expression programs during innate sensing. Nat Immunol. 2016;17:140–9

36. Stielow B, Sapetschnig A, Wink C, Kruger I, Suske G. SUMO-modified Sp3 represses transcription by provoking local heterochromatic gene silencing. EMBO Rep. 2008;9:899–906

37. Tempé D, Vives E, Brockly F, Brooks H, De Rossi S, Piechaczyk M, et al. SUMOylation of the inducible (c-Fos:c-Jun)/AP-1 transcription complex occurs on target promoters to limit transcriptional activation. Oncogene. 2014;33:921–7

38. Cubeñas-Potts C, Matunis MJ. SUMO: A Multifaceted Modifier of Chromatin Structure and Function. Dev Cell. 2013;24:1–12

39. Chymkowitch P, Nguéa PA, Enserink JM. SUMO-regulated transcription: Challenging the dogma. BioEssays News Rev Mol Cell Dev Biol. 2015;37:1095–105

40. Brahimi M, Saidi D, Touhami H, Bekadja MA. The Use of CD45/SSC Dot Plots in the Classification of Acute Leukemias. J Hematol Thromboembolic Dis [Internet]. 2014 [cited 2017 May 2]; Available from: https://www.esciencecentral.org/journals/the-use-of-cdssc-dot-plots-in-the-classification-of-acute-leukemias-2329-8790.1000e107.php?aid=22350

41. Van Parijs L, Refaeli Y, Lord JD, Nelson BH, Abbas AK, Baltimore D. Uncoupling IL-2 signals that regulate T cell proliferation, survival, and Fas-mediated activation-induced cell death. Immunity. 1999;11:281–8

42. Kim YS, Nagy K, Keyser S, Schneekloth JS. Jr. An Electrophoretic Mobility Shift Assay Identifies a Mechanistically Unique Inhibitor of Protein Sumoylation. Chem Biol. 2013;20:604–13

43. Kim YS, Keyser SGL, Schneekloth JS. Jr. Synthesis of 2’,3’,4’-trihydroxyflavone (2-D08), an inhibitor of protein sumoylation. Bioorg Med Chem Lett. 2014;24:1094–7

44. Fukuda I, Ito A, Hirai G, Nishimura S, Kawasaki H, Saitoh H, et al. Ginkgolic acid inhibits protein SUMOylation by blocking formation of the E1-SUMO intermediate. Chem Biol. 2009;16:133–10

45. Hendriks IA, Lyon D, Young C, Jensen LJ, Vertegaal ACO, Nielsen ML. Site-specific mapping of the human SUMO proteome reveals co-modification with phosphorylation. Nat Struct Mol Biol. 2017;24:325–36

46. Subramanian L, Benson MD, Iñiguez-Lluhí JA. A Synergy Control Motif within the Attenuator Domain of CCAAT/Enhancer-binding Protein α Inhibits Transcriptional Synergy through Its PIASy-enhanced Modification by SUMO-1 or SUMO-3. J Biol Chem. 2003;278:9134–41

47. Kim J, Sharma S, Li Y, Cobos E, Palvimo JJ, Williams SC. Repression and Coactivation of CCAAT/Enhancer-binding Protein ε by Sumoylation and Protein Inhibitor of Activated STATx Proteins. J Biol Chem. 2005;280:12246–54

48. Zhou Q, Zhang L, Chen Z, Zhao P, Ma Y, Yang B, et al. Small ubiquitin-related modifier-1 modification regulates all-trans-retinoic acid-induced differentiation via stabilization of retinoic acid receptor α. FEBS J. 2014;281:3032–47

49. Zhu L, Santos NC, Kim KH. Small Ubiquitin-Like Modifier-2 Modification of Retinoic Acid Receptor-α Regulates Its Subcellular Localization and Transcriptional Activity. Endocrinology. 2009;150:5586–95

50. Psakhye I, Jentsch S. Protein Group Modification and Synergy in the SUMO Pathway as Exemplified in DNA Repair. Cell [Internet]. 2012 [cited 2012 Nov 7]; Available from: http://www.sciencedirect.com/science/article/pii/S009286741201241X

51. Ouyang J, Gill G. SUMO engages multiple corepressors to regulate chromatin structure and transcription. Epigenetics. 2009;4:440–4

52. Pascual G, Fong AL, Ogawa S, Gamliel A, Li AC, Perissi V, et al. A SUMOylation-dependent pathway mediates transrepression of inflammatory response genes by PPAR-Y. Nature. 2005;437:759–63

53. Ouyang J, Shi Y, Valin A, Xuan Y, Gill G. Direct binding of CoREST1 to SUMO-2/3 contributes to gene-specific repression by the LSD1/CoREST1/HDAC complex. Mol Cell. 2009;34:145–54

54. Nacerddine K, Lehembre F, Bhaumik M, Artus J, Cohen-Tannoudji M, Babinet C, et al. The SUMO Pathway Is Essential for Nuclear Integrity and Chromosome Segregation in Mice. Dev Cell. 2005;9:769–79

55. He X, Riceberg J, Soucy T, Koenig E, Minissale J, Gallery M, et al. Probing the roles of SUMOylation in cancer cell biology by using a selective SAE inhibitor. Nat Chem Biol. 2017;

